# A photostable genetically encoded voltage indicator for imaging neural activities in tissue and live animals

**DOI:** 10.1101/2025.05.29.656768

**Authors:** Chang Cao, Ruixin Zhu, Shihao Zhou, Zirun Zhao, Chang Lin, Shuzhang Liu, Luxin Peng, Fedor V. Subach, Kiryl D. Piatkevich, Peng Zou

## Abstract

Genetically encoded voltage indicators (GEVIs) enable noninvasive, high-speed monitoring of electrical activity but are constrained by limited brightness and rapid photobleaching under continuous illumination. Here, we present Vega, a highly photostable green fluorescence GEVI with both high sensitivity (ΔF/F = -33% per 100 mV) and fast response (1.34 ms). Under one-photon excitation at 1 W/cm^2^, Vega exhibits more than 20-fold slower photobleaching than the spectrally similar GEVI, Ace-mNeon2. In acute mouse brain slice, Vega enabled wide-field high-fidelity recording of action potentials from 51 neurons simultaneously. In pancreatic islets, it revealed heterogeneous β-cell activation and intercellular coupling in response to glucose elevation. Finally, one-photon imaging in awake mice demonstrated stable cortical voltage mapping *in vivo*. Vega thus overcomes the longstanding photostability-performance trade-off, enabling chronic, high-fidelity voltage imaging across preparations.

## Introduction

Voltage imaging offers noninvasive mapping of membrane-potential dynamics with millisecond temporal and micrometer spatial resolution, thereby surmounting the sampling limitations and invasiveness of electrode recordings in intact neural circuits^1^. Genetically encoded voltage indicators (GEVIs) facilitate cell-type-specific expression and optical readout to track both action potentials and subthreshold events across hundreds to thousands of neurons simultaneously^1-3^. An ideal GEVI should carry high sensitivity (large ΔF/F per 100 mV), ultrafast response kinetics (sub-millisecond on/off rates), and strong photostability, while minimizing phototoxicity to enable prolonged, repeatable imaging sessions with minimal cytotoxicity^4, 5^.

Despite numerous designs, few voltage indicators support a truthful claim of long-term recordings. Rhodopsin-based GEVIs such as Quasar6a deliver high sensitivity (73 ± 8% ΔF/F per 100 mV) but remain intrinsically dim due to low fluorescence quantum yield^6, 7^. One approach to increase brightness is based on enabling electrochromic fluorescence resonance energy transfer (eFRET) between fluorescent proteins (FPs) and voltage-sensing rhodopsins, which are tethered via a short amino acid linker in the protein fusion. While eFRET GEVIs, such as Ace-mNeon2 (−31.3 ± 0.7% ΔF/F per 120 mV) and Cepheid (−33.6 ± 1.5% ΔF/F per 100 mV), are characterized by enhanced brightness, they are limited by the photostability of the FPs, restricting continuous imaging to minutes^2, 3, 8-10^. The ASAP and JEDI series of GEVIs, designed by fusing voltage-sensing domains to circularly permuted GFP (cpGFP), exhibit high voltage sensitivity and sub-millisecond kinetics^11-15^, but undergo rapid, biphasic photobleaching under blue-light illumination, causing up to 30% loss after 5 s at 0.32 W/cm^2^. Alternatively, in hybrid chemigenetic GEVIs, such as HVI and Voltron, FPs are replaced with synthetic dyes characterized by superior photostability, thus extending continuous imaging beyond 15 min, yet they require exogenous dye delivery that complicates chronic experiments *in vivo*^16-20^.

Recent advances in FP engineering have revitalized this field. StayGold, a GFP derived from *Ceratocystis uchidae*, combines high brightness and photostability but has a strong dimerization tendency, precluding its use in protein fusions^21^. Three independent teams have since engineered monomeric StayGold variants^22-24^. Among these, mBaoJin preserves StayGold’s favorable photostability (t_1/2_ = 934 s under 5.1 W/cm^2^ widefield illumination) while maturing rapidly (< 10 min) and exhibiting exceptional chemical and pH stability. mBaoJin also delivers ∼2.5-fold higher intracellular brightness than mNeonGreen. These characteristics establish mBaoJin as an ideal fluorescent moiety to engineer GEVI for extended voltage-imaging experiments, enabling continuous recordings with minimal signal decay^24^.

Herein, we introduce Vega, a green eFRET GEVI that unites exceptional brightness, high sensitivity, and superb photostability for continuous, 15-min recordings. In acute brain slices, Vega enables wide-field imaging of electrical activity from 50 neurons simultaneously with a high signal-to-noise ratio (SNR). In intact pancreatic islets, Vega reveals dynamic intercellular electrical coupling among β cell in response to glucose stimulation. Finally, one-photon wide-field imaging in awake mice demonstrates Vega’s robust *in vivo* performance, allowing chronic monitoring of spontaneous neuronal activity in the primary visual cortex.

## Results

Our group previously developed a red GEVI, Cepheid, by inserting a red fluorescent protein into Ace2 rhodopsin’s first extracellular loop and introducing the D81C mutation to enhance voltage responsiveness^2^. Applying the same design logic, we decided to develop a green eFRET indicator by swapping in the bright, photostable StayGold variant mBaoJin (Figure 1A). To maximize voltage sensitivity, we replaced the original n2 (MVSKGEEEN) and c4 (KGMDELYK) sequences in the N- and C-termini of mBaoJin with the flexible Gly_4_Ser linker.^24^ The resulting Ace2_1-68_-G4S-mBaoJin-G4S-Ace2_69-227_ showed -20.5 ± 0.1% (mean ± SD here and throughout unless otherwise stated) ΔF/F per 100 mV. Truncating this to a single GlySer linker further boosted sensitivity by 1.5-fold (Figure S1). We then replaced mBaoJin with an improved variant, mBaoJin(3M), to extend photostability for chronic imaging. Finally, to enhance neuronal membrane targeting, we appended a Kv2.1-derived soma-targeting motif and an ER-export (ER2) sequence at the C-terminus, yielding Ace2_1-68_-GS-mBaoJin(3M)-GS-Ace2_69-227_-Kv2.1-ER2, termed Vega, for all subsequent experiments (Figure 1A).

**Figure 1.**
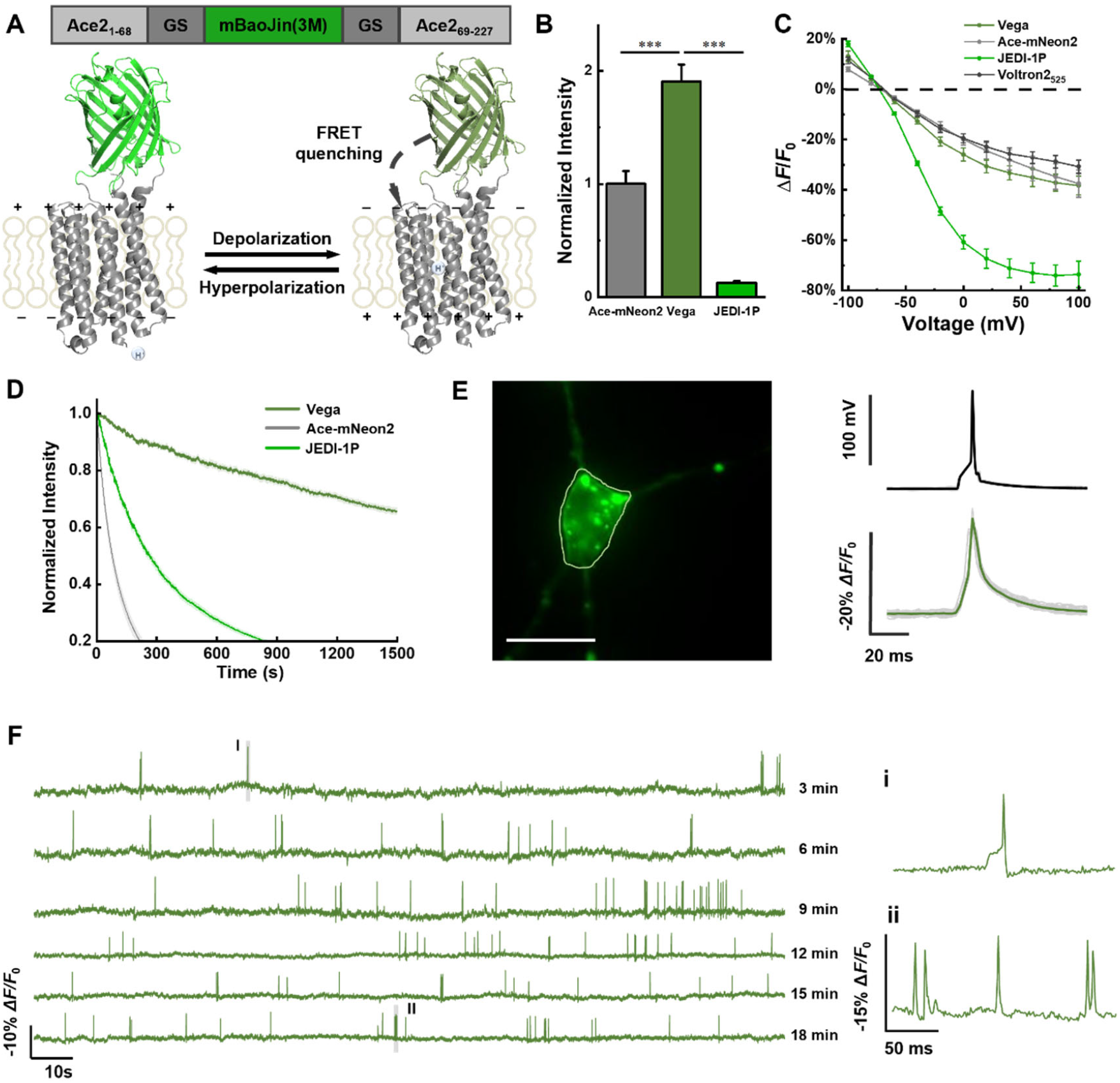
Design and characterization of Vega. **(A)** Structural model of Vega with mBaoJin(3M) inserted into the extracellular loop of Ace2 rhodopsin. **(B)** Normalized whole-cell fluorescence of HEK293T cells expressing Ace-mNeon2 (n = 87 cells), Vega (n = 83 cells) and JEDI-1P (n = 58 cells). Cells were illuminated with 488 nm laser at 3 W/cm^2^. ***P<0.001, one-way ANOVA followed by Bonferroni test. **(C)** F-V curves recorded in HEK cells expressing Vega, Ace-mNeon2, JEDI-1P and Voltron2525. Error bars represent SEM. **(D)** Representative normalized photobleaching curves of Vega, Ace-mNeon2 and JEDI-1P acquired at 10 Hz. The illumination intensity of 488 nm lasers was 3 W/cm^2^. **(E)** Optical waveforms of Vega to APs. Region of interest that yielded the trace is marked by green circle. Scale bars, 10 μm. **(F)** Six rounds of 3-min voltage imaging at 400 Hz were performed successively for cultured neurons. The illumination intensity of 488 nm laser was 1 W/cm^2^. Two zoomed-in signals (i-ii) from two shaded regions (I-II) were presented on the left.

When expressed in human embryonic kidney 293T (HEK293T) cells, Vega was ∼2-fold brighter than the eFRET sensor Ace-mNeon2 and ∼8-fold brighter than JEDI-1P under 488 nm excitation at 3 W/cm^2^ (Figure 1B). We applied whole-cell voltage clamp to vary the membrane potential from -70 mV to 30 mV, while simultaneously recording the Vega fluorescence. Vega demonstrated greater sensitivity (ΔF/F = -33.2 ± 2.8% per 100 mV) than both Ace-mNeon2 (−26.7 ± 3.2%) and Voltron2 (−27.5 ± 2.1%) (Figure 1C). While its voltage sensitivity was lower than that of JEDI-1P (−66.3 ± 2.5%), Vega exhibited comparably fast kinetics (τ_on_ = 1.34 ms, τ_off_ = 3.74 ms) relative to JEDI-1P (τ_on_ = 5.41 ms, τ_off_ = 2.91 ms) at 21–23 °C (Figure S2) ^15^. Under continuous 488 nm illumination (1 W/cm^2^), Vega’s fluorescence decayed by only 25% after ∼769 ± 297 s (t_0.75_), exceeding Ace-mNeon2’s t_0.75_ of 38 ± 5 s by >20-fold and JEDI-1P’s t_0.75_ of 102 ± 21 s by >7.5-fold (Figure 1D).

In cultured hippocampal neurons, Vega reports single action potentials with a ΔF/F of -17.1 ± 2.2% per spike (whole cell fluorescence, n = 5 neurons), surpassing Ace-mNeon2’s -13.2 ± 1.4% per spike (n = 4 neurons) under identical patch-clamp protocols (Figure 1E and S3). However, Vega exhibits intracellular puncta, indicating suboptimal membrane trafficking that may be alleviated by additional export motifs (Figure 1E). During 18 min of continuous 488 nm, 1 W/cm^2^ illumination, Vega’s SNR declines by <10% over 15 min (Figure 1F), significantly more photostable than the chemigenetic indicator Voltron2 under similar conditions (41% over 10 min)^20^. Neuron morphology and spontaneous firing rates remain statistically unchanged after six imaging rounds (Figure S4), confirming negligible phototoxicity or functional perturbation. Vega reliably resolves subthreshold events, such as depolarizations and repolarizations, throughout prolonged recordings (Figure 1F and Figure S5), and its baseline fluorescence bleaches at ∼1.5% per min, on par with the most photostable opsin-based GEVIs under widefield illumination.

Because Vega reliably reports both tonic and burst action potentials as well as subthreshold depolarizations with a high SNR during prolonged imaging sessions, we exploited its superior photostability to monitor drug-induced modulation of neuronal excitability in cultured rat hippocampal neurons. In DIV 14 neurons expressing Vega, we applied cyclothiazide (CTZ), a potent positive allosteric modulator of AMPA receptors that prevents receptor desensitization and amplifies glutamatergic currents, to induce epileptiform discharges^25^. CTZ increased the mean spontaneous firing rate from 0.12 ± 0.06 Hz at baseline to 0.22 ± 0.08 Hz at 2 μM, and further to 1.84 ± 0.66 Hz at 10 μM, demonstrating a clear dose-dependent enhancement of excitability^26^ (Figure 2A-B).

**Figure 2.**
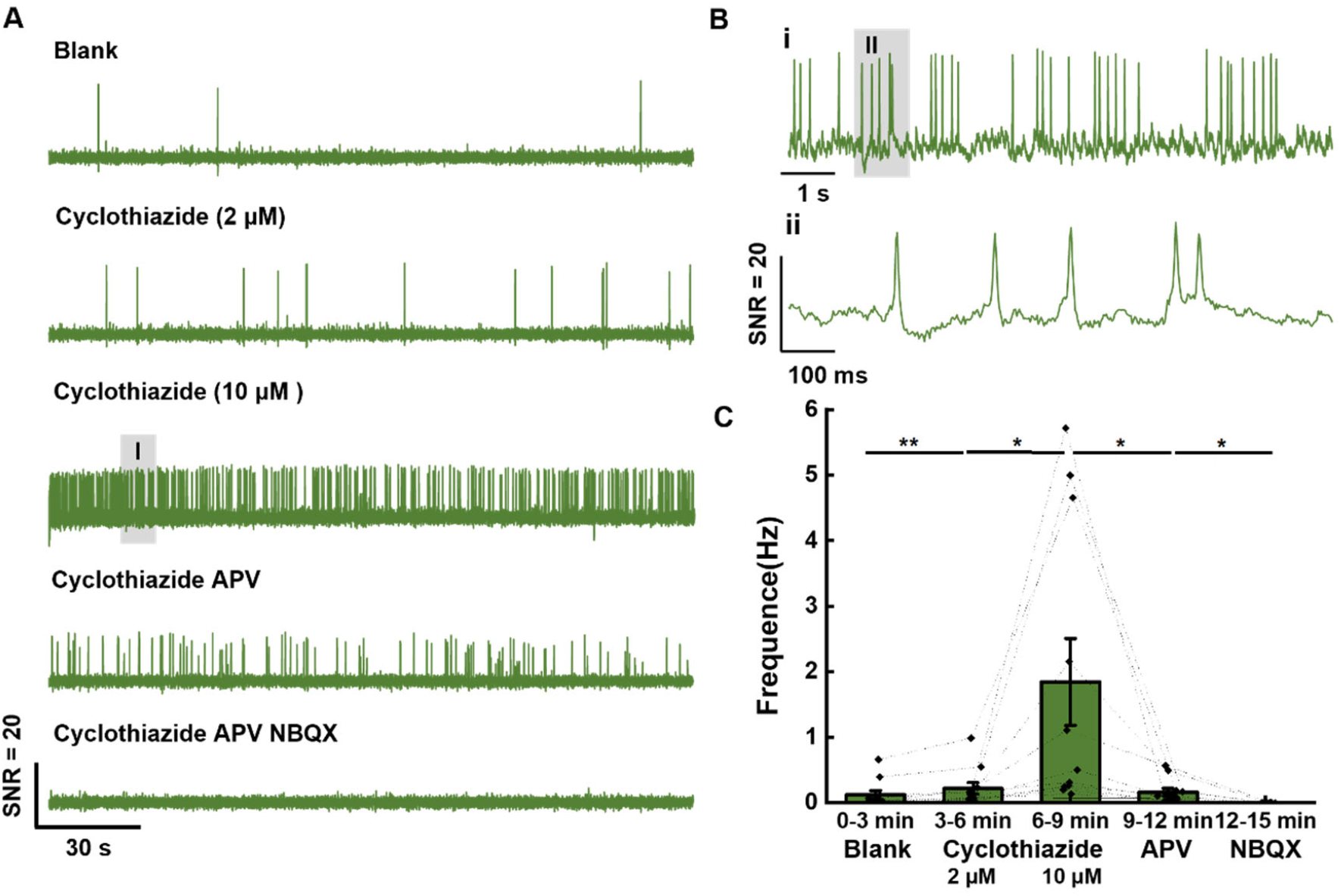
Recording of drug-induced excitability changes in cultured neuron. **(A)** Five rounds of 3-min voltage imaging in cultured neuron treated with different drugs: 2 and 10 μM cyclothiazide was added before the 2^nd^ and 3^rd^ round, respectively, followed by 25 μM APV and 10 μM NBQX in the last two rounds, respectively. The fluorescence signal of Vega was corrected for photobleaching. The illumination intensity of the 488 nm laser was 1 W/cm^2^, and the camera frame rate was 400 Hz. **(B)** Two zoomed-in signals (i-ii) from two shaded regions (I-II). **(C)** Statistics of AP frequency during each round of recording with drug treatment (n = 7 cells). Statistical significances are determined by paired t-test (* p < 0.05). Error bars represent SD.

To parse the receptor contributions to hyperexcitability, we replaced CTZ with two selective glutamate receptor antagonists. Incubating neurons in D-APV (50 µM), a competitive NMDA receptor antagonist, reduced spontaneous firing from 1.84 ± 0.66 Hz to 0.16 ± 0.06 Hz, an approximately 11.5-fold suppression consistent with blockade of NMDA-dependent EPSP components (Figure 2C). In contrast, NBQX (10 µM), a potent AMPA/KA receptor antagonist, completely abolished synaptic firing, effectively silencing the network (Figure 2C). Measuring the same neuron across treatment conditions eliminates intercellular variability and enhances the statistical power of excitability assays. These data demonstrate that Vega can continuously monitor action potential firing under distinct pharmacological manipulations, offering an all-optical alternative to patch-clamp recordings.

Next, we evaluated Vega performance for voltage imaging in acute brain slices. We injected adeno-associated virus (AAV) encoding Vega into the mouse brain and prepared acute slices three weeks later. Confocal imaging of formaldehyde-fixed tissue confirmed strong Vega expression in the cortex and hippocampus (Figure S6). Importantly, whereas cultured hippocampal neurons exhibited intracellular puncta, Vega in brain slices localized predominantly to the plasma membrane.

We recorded spontaneous signals from thalamic neurons. Vega supported continuous imaging sessions, up to 15 minutes at a 400 Hz camera frame rate, while maintaining high sensitivity (−12 ± 1.4% ΔF/F) and SNR (12.6 ± 2.2) per AP (Figure 3A). Over the course of imaging, Vega’s SNR remained high with no significant drop, underscoring its exceptional photostability (Figure 3B), and we observed no obvious changes in the full-width at half-maxima (FWHM) of optically recorded AP waveforms (Figure 3C). By contrast, Ace-mNeon2 showed 30% decline in SNR over 7-min imaging session under similar conditions. (Figure S7).

**Figure 3.**
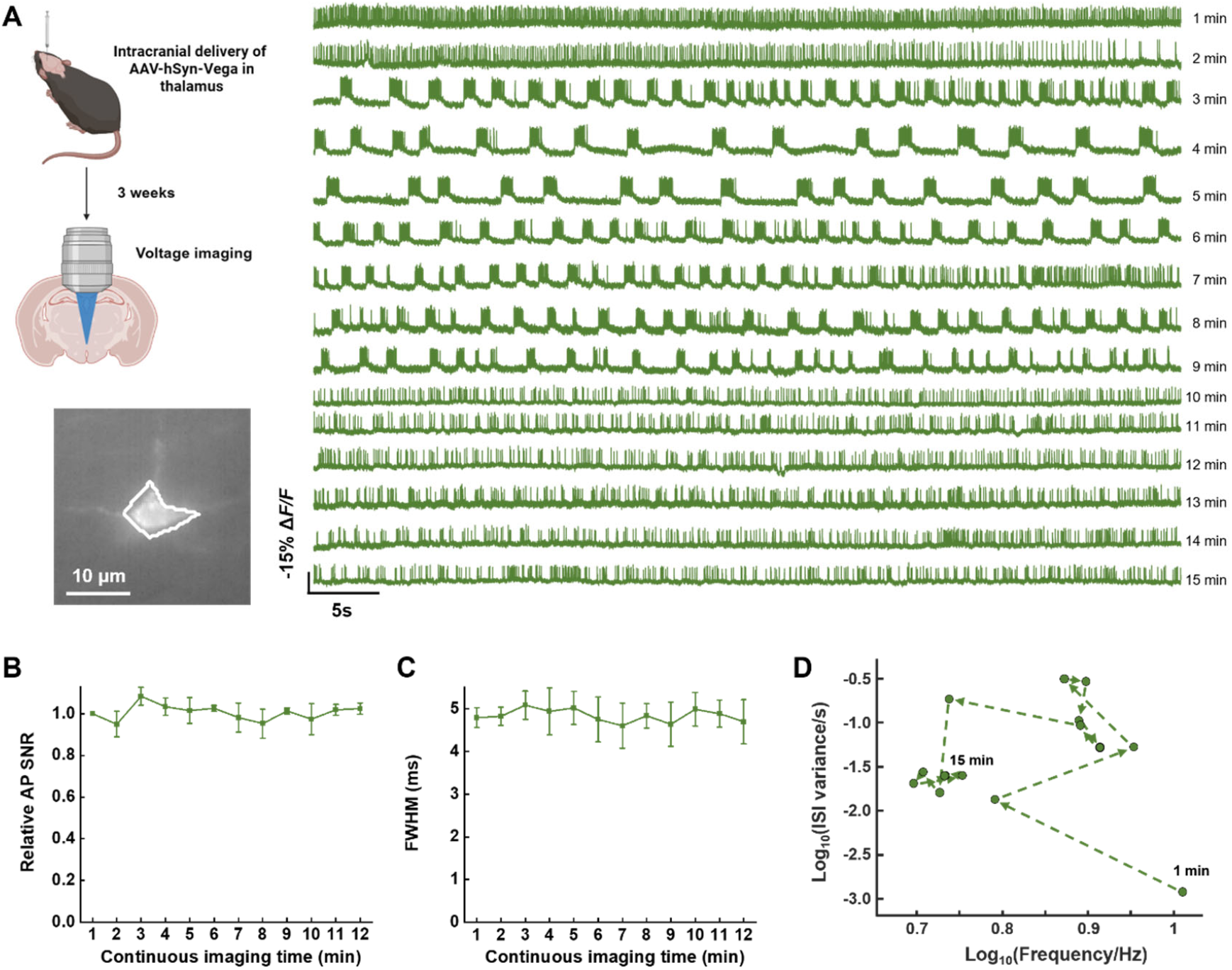
Voltage imaging with Vega in acute brain slices. **(A)** Fluorescence imaging of a thalamic neuron expressing Vega in acute brain slice. Left: schematic of AAV injection and wide-field fluorescence image of a neuron expressing Vega in the thalamus. Scale bar, 10 μm. Right: whole-cell fluorescence (outlined by the polygon) of the neuron shown on the left, recorded at 400 Hz under 488 nm laser illumination at 1 W/cm^2^. **(B-C)** Normalized SNR (B) and averaged FWHM (C) of APs recorded by Vega (n = 3 cells in 3 mice) in each 1-min imaging session. **(D)** Plot of the variance of inter-spike-intervals (ISI) against AP frequencies in log scale of the neuron in **(A)**. Arrows indicate the sequence from 1 to 15 min.

We observed reversible shifts in neuronal firing patterns between tonic and burst modes during long-term imaging. For each 1-min imaging session, we quantified the AP frequency and the variance of inter-spike-interval (ISI). Burst activity was characterized by a high variance in ISI. A scatter plot of ISI variance versus AP frequency reveals the trajectory of firing pattern shift over time (Figure 3D). Data analysis revealed three distinct clusters of data points based on characteristics of ISI and frequency, with each cluster exhibiting temporal segregation. Because Vega exhibits low phototoxicity, these phase-dependent shifts reflect true neuronal state changes rather than imaging artifacts. This result demonstrates how extended voltage imaging can uncover dynamic modulations in neuronal signaling and functional state transitions.

To achieve wide-field voltage imaging with Vega, we built a customized microscope equipped with a 25x water-immersion objective (Nikon) and a 60-mm tube lens, achieving an effective magnification of 6.5x. Using the central 512-by-512 pixels of a Hamamatsu ORCA-Fusion sCMOS camera, a 512 um by 512 um field of view of the brain slice sample was imaged at 400 Hz camera frame rate (Figure 4A). A total of 51 thalamic neurons were identified and recorded with high SNR (16.3 ± 5.6), revealing heterogeneous firing patterns across the population (Figure 4B). While approximately one third of the neurons fired tonically during the 2-min imaging session, the remaining exhibited burst firing patterns with ISI varying from less than 10 seconds to longer than 1 minute. Notably, high-frequency spiking neurons (>1 Hz) predominantly displayed tonic firing patterns. Consistent with this observation, more quantitative analysis revealed that, on a logarithmic scale, spike frequency and ISI variance were strongly, negatively correlated (R^2^ = 0.76, p < 0.05) (Figure 4C).

**Figure 4.**
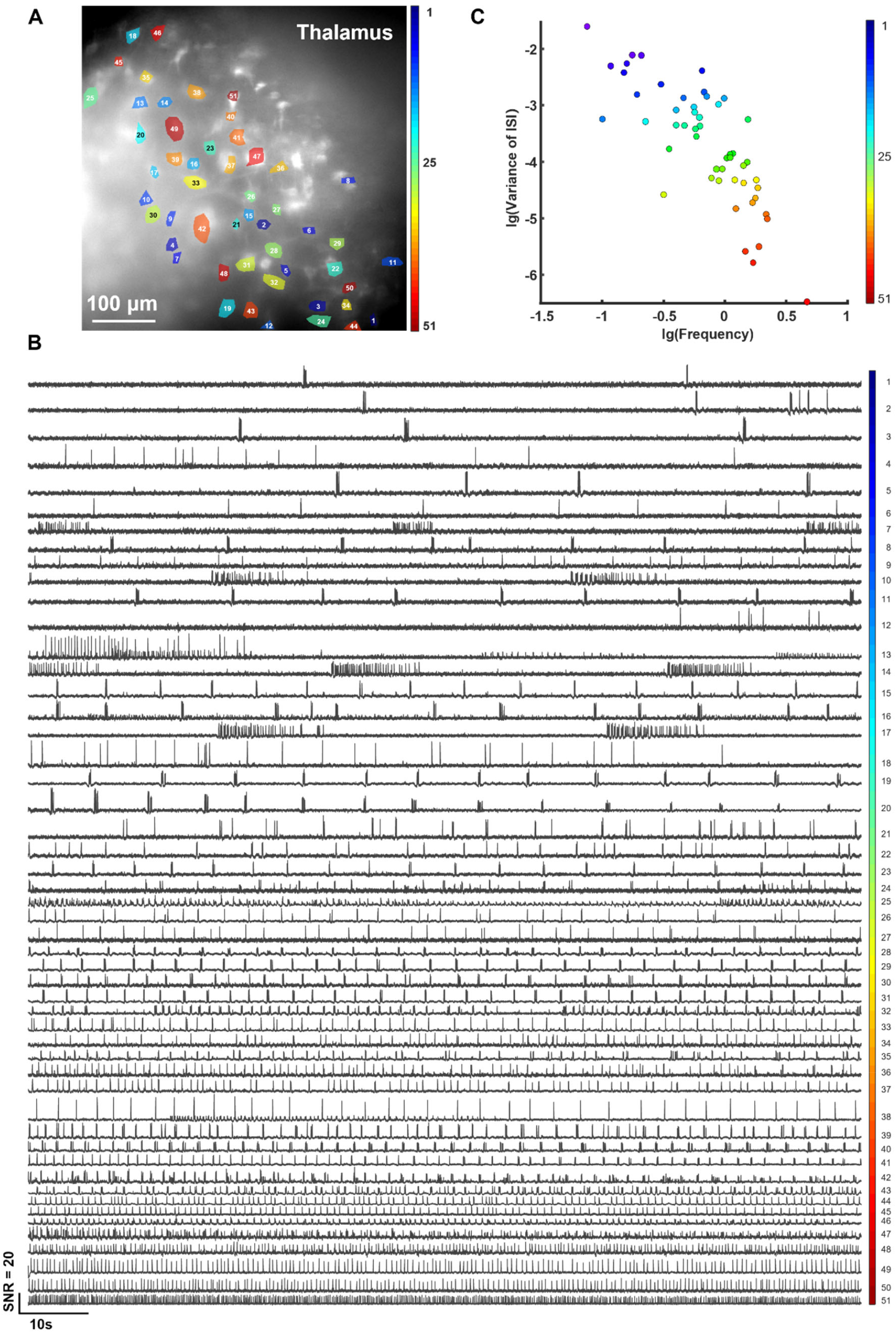
Wide-field voltage imaging of multiple neurons with Vega simultaneously in acute brain slice. **(A)** Representative image of acute brain slice expressing Vega in the thalamus. Scale bar, 100 μm. **(B)** SNR traces indicating spontaneous activity from the labeled neurons in **(A)**. The illumination intensity of 488 nm laser was 1.5 W/cm^2^. The camera frame rate was 400 Hz. **(C)** lg(Variance of inter-spike-intervals) plotted against lg(Frequency) between neurons labelled in **(A)**.

We assigned each neuron an index based on its orthogonal projection onto the linear fit. As the index rose, our pseudocolor map shifted from blue to red, marking a gradual transition from burst-dominated to tonic firing. To test whether similarly indexed neurons cluster spatially, we mapped examples (indices 7, 10, 14, 17) and found no localized grouping, with burst- and tonic-pattern neurons intermingled throughout the slice (Figure 4A). This dispersion suggests that firing-mode diversity likely arises from intrinsic cellular or circuit properties rather than anatomical sub-regions.

We next used Vega to visualize glucose-induced electrical activity in intact pancreatic islets. When glucose levels rise, Glut2 transporters, K_ATP_ channels, and voltage-gated Ca^2+^ channels coordinate to depolarize β cell membranes within one minute^27, 28^. Because neighboring β cells are electrically coupled via gap junctions, this depolarization propagates as synchronized waves, producing spatially coordinated voltage dynamics across the islet^29, 30^. Although patch-clamp studies have characterized single-cell electrical responses^31-33^, the lack of long-term, high-throughput imaging methods for intact islets has limited our insight into these multicellular spatial patterns.

To achieve β cell-specific expression of Vega, isolated mouse islets were infected with an adenovirus encoding prip2::Vega and imaged ex vivo 36-48 hours later (Figure 5A). To trigger electrical activity, islets were first held in 3 mM glucose (suppressing native action potentials), then perfused with 13 mM glucose during imaging. Over a continuous 15-min recording, Vega faithfully tracked membrane potential changes in β cells, capturing both action potentials and subthreshold depolarizations (Figure 5B). Minimal photobleaching of the fluorescence baseline enabled direct measurement of depolarization and repolarization kinetics from the raw traces.

**Figure 5.**
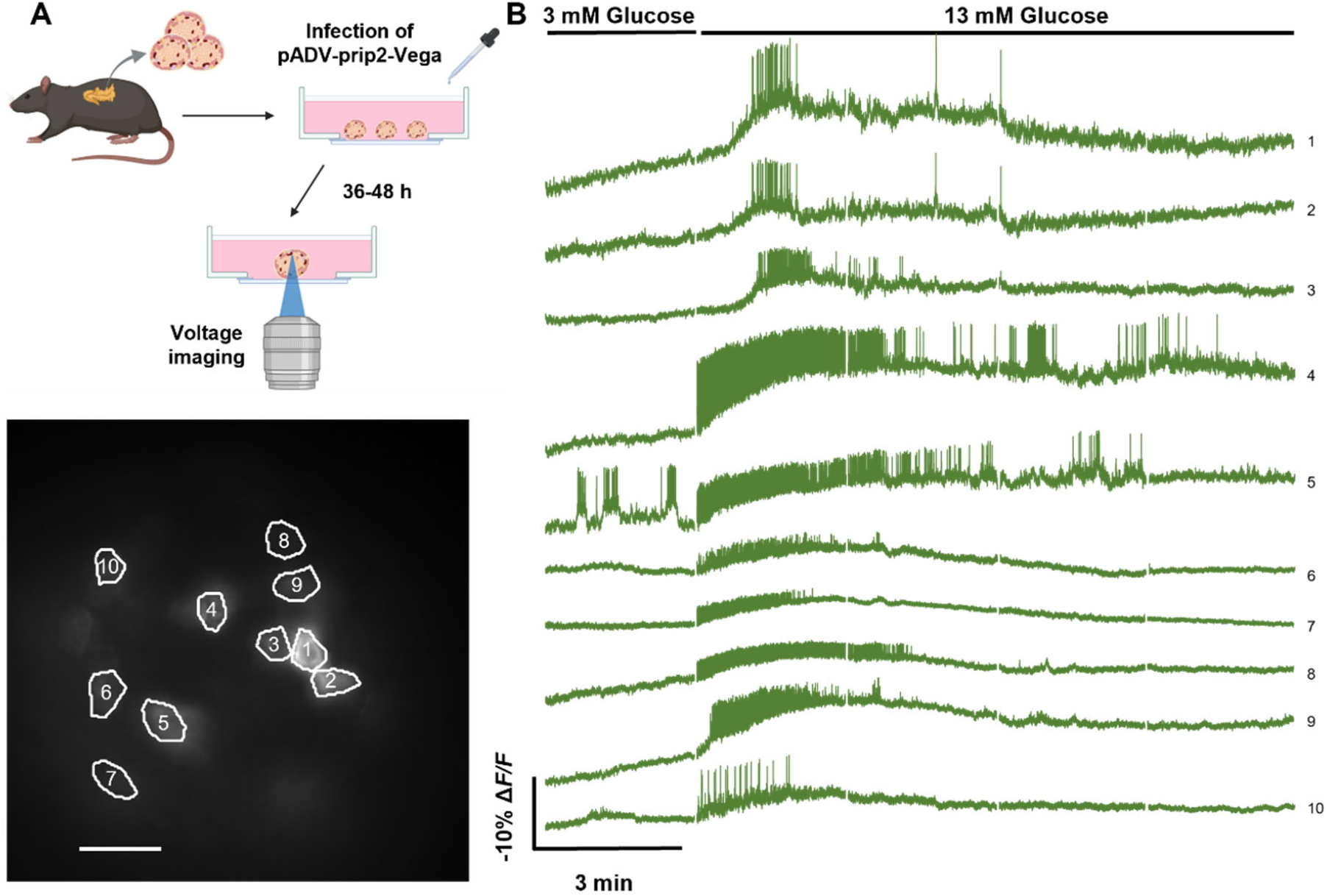
Vega enables 15-minute long-term imaging of mouse pancreatic islets. **(A)** Schematic of virus infection and setup of imaging experiment in mouse pancreatic islets. Representative image of β cells expressing Vega in mouse pancreatic islet is shown at the bottom. Scale bar, 100 μm. **(B)** Five rounds of 3-min voltage imaging successive in mouse pancreatic islets. 13 mM glucose was replaced with 13 mM glucose after imaging for 3 minutes. The numbers on right correspond to cells marked in **(A)**. Raw traces were shown without photobleaching correction. The illumination intensity of 488 nm laser was 1.5 W/cm^2^ and the camera frame rate was 200 Hz.

Under low glucose (3 mM), nearly all β cells were electrically quiescent. Upon stimulation with 13 mM glucose, their voltage traces diverged markedly, reflecting population heterogeneity in activation dynamics. Some cells (e.g., #1-3) underwent a gradual depolarization followed by rhythmic action-potential bursts, whereas others (e.g., #4-5) activated more rapidly, firing at higher frequencies with sustained oscillatory bursting after 3-4 min. A third group (e.g., #6-10) produced a brief burst and then slowly repolarized. Closer inspection of raw fluorescence traces revealed local synchronization events. For instance, simultaneous bursts were observed in adjacent cell pairs (e.g., cells 1/2 and 8/9 in Figure 5C), consistent with gap-junction coupling reported in intact islets^29-31^. These findings highlight intrinsic functional diversity among β cells during glucose-induced excitation, underscoring the value of continuous, high-resolution voltage imaging for revealing cell-specific dynamics.

Finally, we evaluated Vega’s performance in awake mice by imaging activities in layer 2/3 pyramidal neurons of the primary visual cortex. We drove Vega expression via AAV infection and recorded spontaneous activity via one-photon epifluorescence microscopy (Figure 6A). Under LED illumination at ∼5 W/cm^2^, typical for imaging green GEVIs *in vivo*^3^, Vega reliably resolved spikes from 9 neurons across a 104 μm-by-156 μm field of view (Figure 6C). In the meantime, we found that Vega can be imaged over 15 minutes with minimal drop in SNR (Figure S8). Although Vega’s SNR (4.0-5.6) is slightly lower than Voltron2’s (∼7) under similar experimental conditions (∼1.8 W/cm^2^)^20^, Vega is fully genetically encoded and requires no exogenous dyes, which greatly simplifies *in vivo* applications. Thus, Vega is compatible with standard, cost-effective optical setups and well suited for extended, longitudinal voltage imaging in the mouse brain.

**Figure 6.**
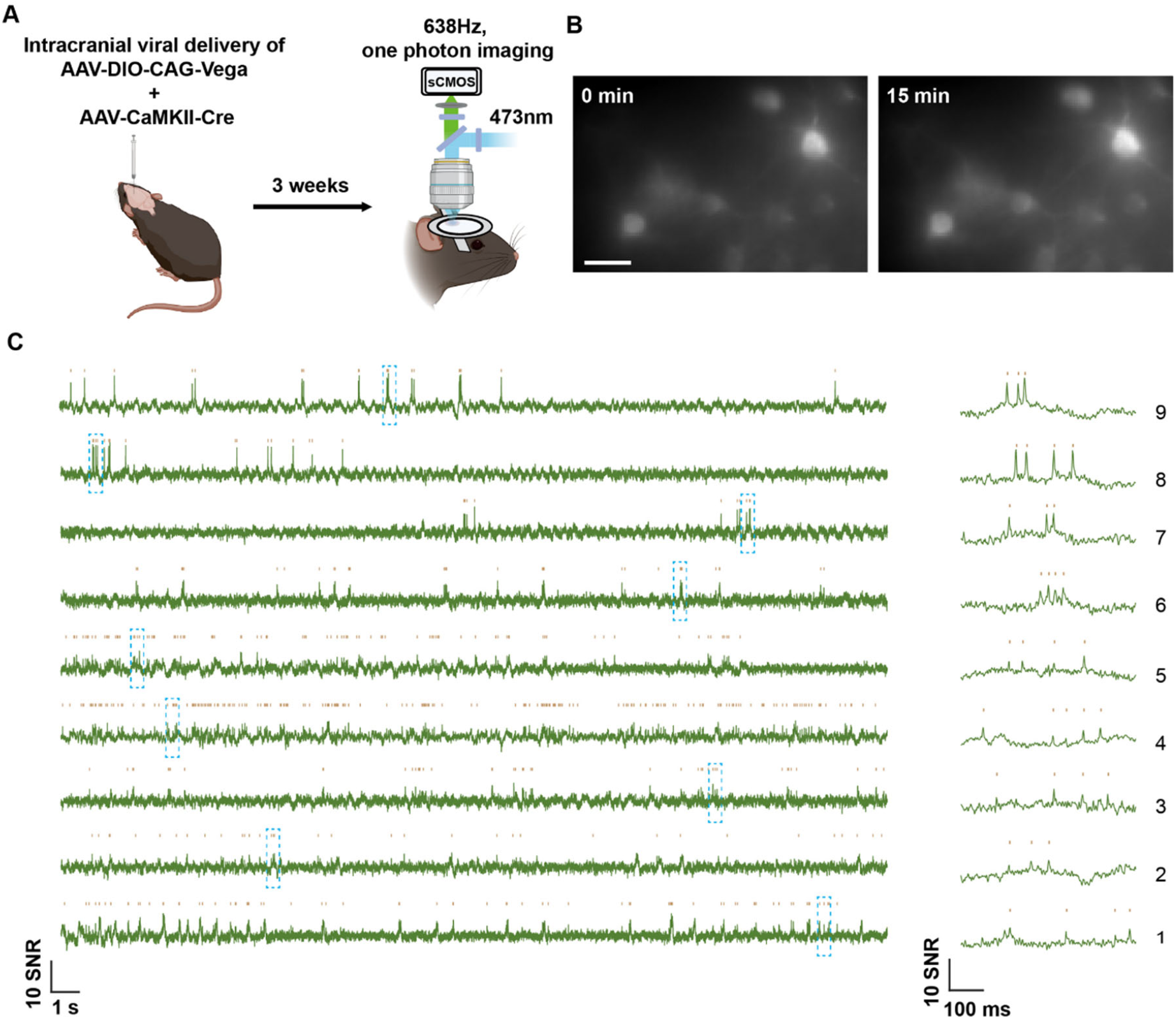
One-photon voltage imaging of Vega expressed in V1 layer 2/3 pyramidal neurons (PNs). **(A)** Schematics of AAV injection and setup of imaging experiment in awake mice. **(B)** Example projection image showing a FOV of L2/3 PNs in V1 expressing Vega *in vivo* for both the beginning and 15-min timepoint. **(C)** 30-sec trace recorded from PNs in L2/3 V1. A 500 ms trace snippets were shown on the right and highlighted in cyan dashed box in the 30-sec recording. Scale bar = 25μm. n = 9 neurons recorded from 2 mice.

## Discussions

Developing a photostable GEVI remains a critical challenge in neuronal electrophysiology. Many neural activities, from circadian rhythms to awake-behavior circuits, unfold over minutes to hours and demand continuous monitoring^34, 35^. GEVIs developed to date have suffered photobleaching that precludes such long-term imaging, forcing researchers to choose between low-throughput electrophysiology or calcium indicators that miss fast and subthreshold events. Here we present Vega, a green GEVI engineered to overcome the above trade-off. Vega’s exceptional photostability supports uninterrupted voltage imaging with high spatial and temporal resolution in both *ex vivo* brain tissue and *in vivo* preparations.

In cultured neurons, Vega’s sustained performance allows systematic testing of pharmacological agents on neuronal excitability, opening the door to long-term, high-throughput voltage assays. In acute brain slice from mouse thalamus, simultaneous recordings from dozens of neurons revealed pronounced heterogeneity in firing patterns among neighboring cells. In mouse pancreatic islets, we tracked β cell responses during glucose elevation, observing both diverse individual dynamics and clear electrical coupling between adjacent cells, findings that align with prior patch-clamp studies. Looking ahead, pairing Vega with red-shifted calcium or voltage indicators will enable multiparametric mapping of intercellular electrical correlations via genetic labeling^2, 3, 36^.

Membrane targeting is essential for the development of any GEVI. Only probes at the plasma membrane translate voltage changes into fluorescence, whereas cytosolic aggregates merely add background noise. In cultured neurons, Vega shows some intracellular aggregation (Figure 1E), which results in reduced sensitivity and SNR relative to HEK293T recordings. However, when expressed via AAV in acute slices, membrane localization is dramatically improved (Figure S6), restoring SNR sufficient for unambiguous spike detection. The improved trafficking *in vivo* might be attributed to lower expression levels achieved via viral delivery.

Finally, Vega’s performance in wide-field, one-photon imaging of the awake mouse primary visual cortex demonstrates its potential to perform longitudinal imaging *in vivo*. However, because one-photon excitation penetrates only superficial cortical layers, deeper recordings demand two-photon voltage imaging to improve both imaging depth and SNR^37^. Recent advances in high-speed, scanning two-photon microscopy have made it possible to record from head-fixed mice with the VSD-based indicator JEDI-2P^12, 14^. Moreover, two-photon excitation at higher illumination power has been shown to enable *in vivo* imaging of the rhodopsin-based GEVI Voltron2, establishing the feasibility of rhodopsin-based indicators under two-photon conditions^38^. Under non-scanning two-photon illumination, rhodopsin-based probes also exhibit greater resting brightness than JEDI-2P, indicating potential for even higher SNR^39, 40^. Collectively, these findings underscore Vega’s promise for two-photon applications.

In the parallel study, Zhang et al. designed a set of bright and photostable GEVIs by swapping mNeonGreen FP in pAce and Ace-mNeon2 with the enhanced mBaoJin variant. Similarly to Vega, integration of mBaoJin variant results in enhanced brightness and photostability of Electras, although with some reduction of voltage sensitivity compared to the mNeonGreen-based counterpart. We did not compare the performance of Vega and Electra side-by-side.

Given Vega’s exceptional brightness and photostability, we anticipate it will resist voltage-sensitivity loss, sustaining SNR under intense illumination, and extend the temporal window for deep-tissue imaging. Moving forward, future efforts would focus on engineering a Vega-based two-photon voltage indicator and optimizing illumination strategies to achieve high-SNR, long-term recordings of neuronal activity *in vivo*. Coupled with advances in high-speed *in vivo* microscopy, Vega is poised to revolutionize large-scale functional mapping of neuronal populations in behaving animals.

## Acknowledgments

We thank Hanbin Zhang from Westlake University for help with the cloning and AAV production. This work was supported by the Ministry of Science and Technology (2022YFA1304700), the National Natural Science Foundation of China (32088101 to P.Z., 32171093 to K.D.P.), and Beijing National Laboratory for Molecular Sciences (BNLMS-CXXM-202403). This work was also in part supported by the Foundation of Westlake University, Westlake Laboratory of Life Sciences and Biomedicine, and ‘Pioneer’ and ‘Leading Goose’ R&D Program of Zhejiang (grant no. 2024SSYS0031 to K.D.P.) and carried out within the state assignment of NRC “Kurchatov Institute” to F.V.S.

## Author Contributions

C.C. and P.Z. conceived the project and designed experiments; C.C., R.Z., Z.Z., C.L. and S.L. performed experiments; F.V.S. engineered mBaoJin(3M). S.Z. performed one-photon imaging in mouse *in vivo*. C.C., S.Z., L.P. and P.Z. analyzed data; C.C., S.Z., K.D.P., and P.Z. wrote the paper with inputs from all authors.

