## Supplementary information for "A photostable genetically encoded voltage indicator for imaging neural activities in tissue and live animals"

### Supplementary Tables

#### Table S1. Photobleaching quarter-life of Vega and Ace-mNeon2

| Voltage indicators | Excitation wavelength (nm) | Illumination intensity ($W \cdot cm^{-2}$) | Emission filter (nm) | Photobleaching $t_{0.75}$ (s) [a] |
|---|---|---|---|---|
| Vega | 488 | 3.0 | 525 / 50 | 769 ± 297, n = 14 |
| Ace-mNeon2 | 488 | 3.0 | 525 / 50 | 38 ± 5, n = 12 |
| JEDI-1P | 488 | 3.0 | 525 / 50 | 102 ± 21, n = 13 |

[a] measured in live HEK293T cells expressing Vega, Ace-mNeon2 and JEDI-1P. "±" represents s.d.

#### Table S2. Voltage sensitivities of Vega and Ace-mNeon2

| Voltage indicators | $\Delta F/F_0$ per AP (%) [a] | SNR [a] | $\Delta F/F_0$ per 100 mV (%) [b] |
|---|---|---|---|
| Vega | -17.1 ± 2.2, n = 5 | 83.7 ± 5.8 | -33.1 ± 2.7, n = 6 |
| Ace-mNeon2 | -13.2 ± 1.4, n = 4 | 80.2 ± 17.4 | -26.7 ± 3.2, n = 8 |

[a] measured in cultured rat neurons at 484 Hz under the illumination of 488 nm (3.0 $W \cdot cm^{-2}$).

[b] measured in HEK293T cells at 1058 Hz under the illumination of 488 nm (3.0 $W \cdot cm^{-2}$).

"±" represents s.e.m.

#### Table S3. Imaging apparatus for fluorescent imaging

| Indicators | Fluorophores | Excitation wavelength (nm) | Emission filter (nm) |
|---|---|---|---|
| Vega | mBaoJin(3M) | 488 | 525 / 50 |
| Ace-mNeon2 | mNeonGreen | 488 | 525 / 50 |
| JEDI-1P | cpGFP | 488 | 525 / 50 |
| Voltron2 | JF525 | 525 | 585 / 65 |

Dichroic mirror: ZT405/488/532/642rpc

**Table S4. List of reagents used in this study**

| Reagent | Vendor | Catalog Number |
|---|---|---|
| DNA extraction kit | TIANGEN | DP118-02 |
| Dulbecco's Modified Eagle's medium (DMEM) | Gibco | C11995500BT |
| Fetal Bovine Serum (FBS) | Gibco | 100099044 |
| Trypsin-EDTA (0.25%) | Gibco | 25200056 |
| Neurobasal™ Medium | Gibco | 21103049 |
| B-27™ Supplement | Gibco | 17504044 |
| GlutaMAX™ Supplement | Gibco | 35050061 |
| Pancreatin | Sigma | P3292 |
| Penicillin-streptomycin | Beyotime | C0222 |
| Tyrode’s Salts Solution | macgene | CC018 |
| Adenosine 5’-triphosphate disodium (ATP·$Na_2$) | Amresco | 0220 |
| Guanosine-5'-triphosphate disodium (GTP·$Na_2$) | Yuanye | 56001-37-7 |
| Magnesium acetate | Sinopharm | 30110518 |
| Matrigel® Matrix | Corning | 356234 |
| poly-D-lysine | Sigma | P7280-5X5M |
| Laminin Mouse Protein | Gibco | 23017015 |
| Opti-MEM™ Medium | Gibco | 31985062 |
| Lipofectamine™ 2000 Reagent | Invitrogen | 11668019 |
| Lipofectamine™ LTX & PLUS™ | Invitrogen | 15338100 |
| Formaldehyde solution | Aladdin | F111934 |
| HEPES | Amresco | 0511 |
| EGTA | Sigma | 03777-10G |
| 2-APB | Abcam | ab120124 |
| Gabazine | Abcam | ab120042 |
| NBQX | Abcam | ab120045 |
| D-AP5 (APV) | Abcam | ab120003 |
| Cyclothiazide | Abcam | ab120061 |

### Supplementary Figures

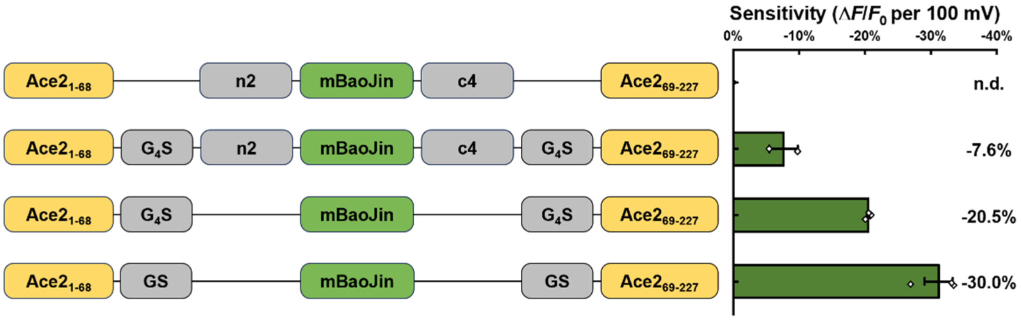

**Figure S1. Optimization of struction in HEK293T cells.** The constructs that have been screened through patch clamp and the screening results in HEK293T cells. Average sensitivity of each construct was presented on the bar, and the sensitivity of mBJ-ΔnΔc-TS was not detected due to poor membrane localization. Error bars represent SEM. $G_4S$: GGGGS, n2: MVSKGEEEN, c4: KGMDELYK.

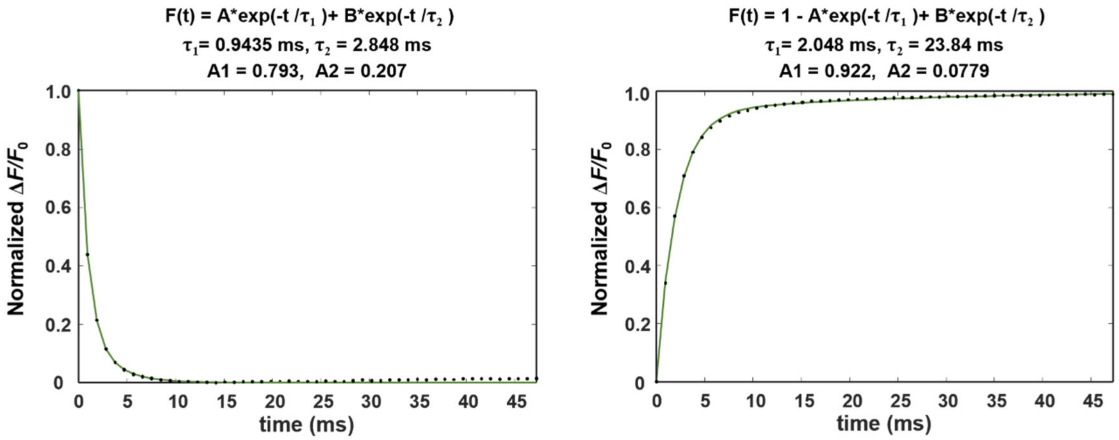

**Figure S2. Kinetic analysis of depolarization and repolarization of Vega.** Step responses of Vega to +100 mV command voltage steps (from -70 mV to +30 mV), normalized to maximum $\Delta F/F_0$ response to the command voltage. The dot are the average values of traces tested in 6 HEK293T cells, and the green line represents the result obtained from a double-exponential fitting.

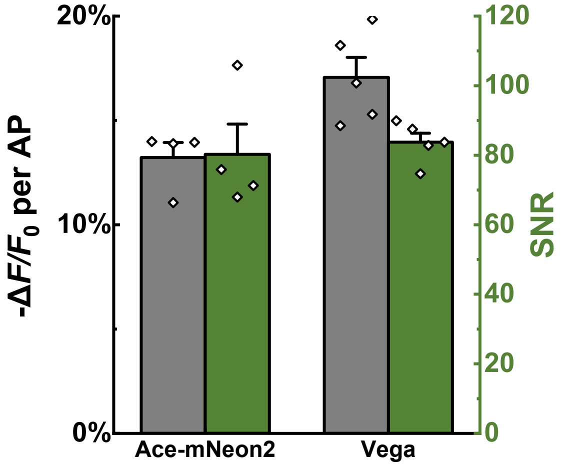

**Figure S3. Comparison of the sensitivity and SNR of Vega with Ace-mNeon2 in cultured neurons.** Neurons expressing Ace-mNeon2 (n = 4 cells) or Vega (n = 5 cells). Error bars represent s.d.

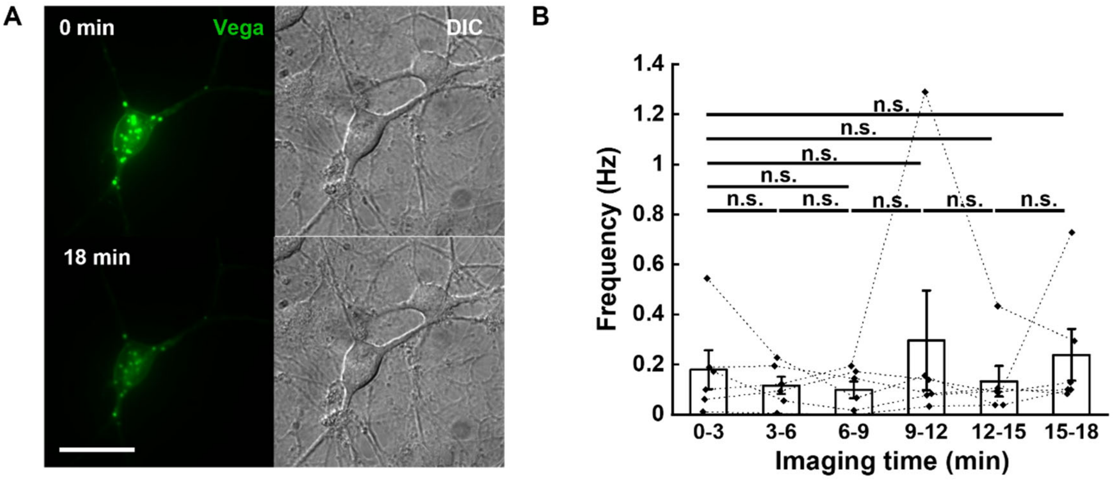

**Figure S4. Vega enables 15-min long-term voltage imaging in cultured neurons. (A)** Wide-field imaging of neuron expressing Vega before/after continuous illumination, scale bar = 50 μm. **(B)** Statistics of AP burst frequency in cultured neurons during six rounds of 3-min voltage imaging. Imaging was performed for each neuron expressing Vega (n = 6 cells). Statistical significances are determined by One-Way Repeated Measures ANOVA. Error bars represent s.e.m.

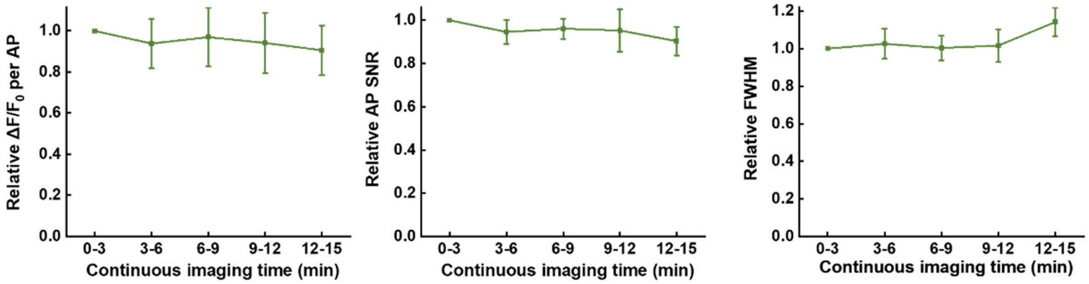

**Figure S5. Statistics of APs of Vega during long-term continuous imaging in cultured neurons.** Normalized sensitivity (left), SNR (middle) and FWHM (right) of APs recorded by Vega (n = 5 cells) in each 3-min imaging session. Error bars represent s.e.m.

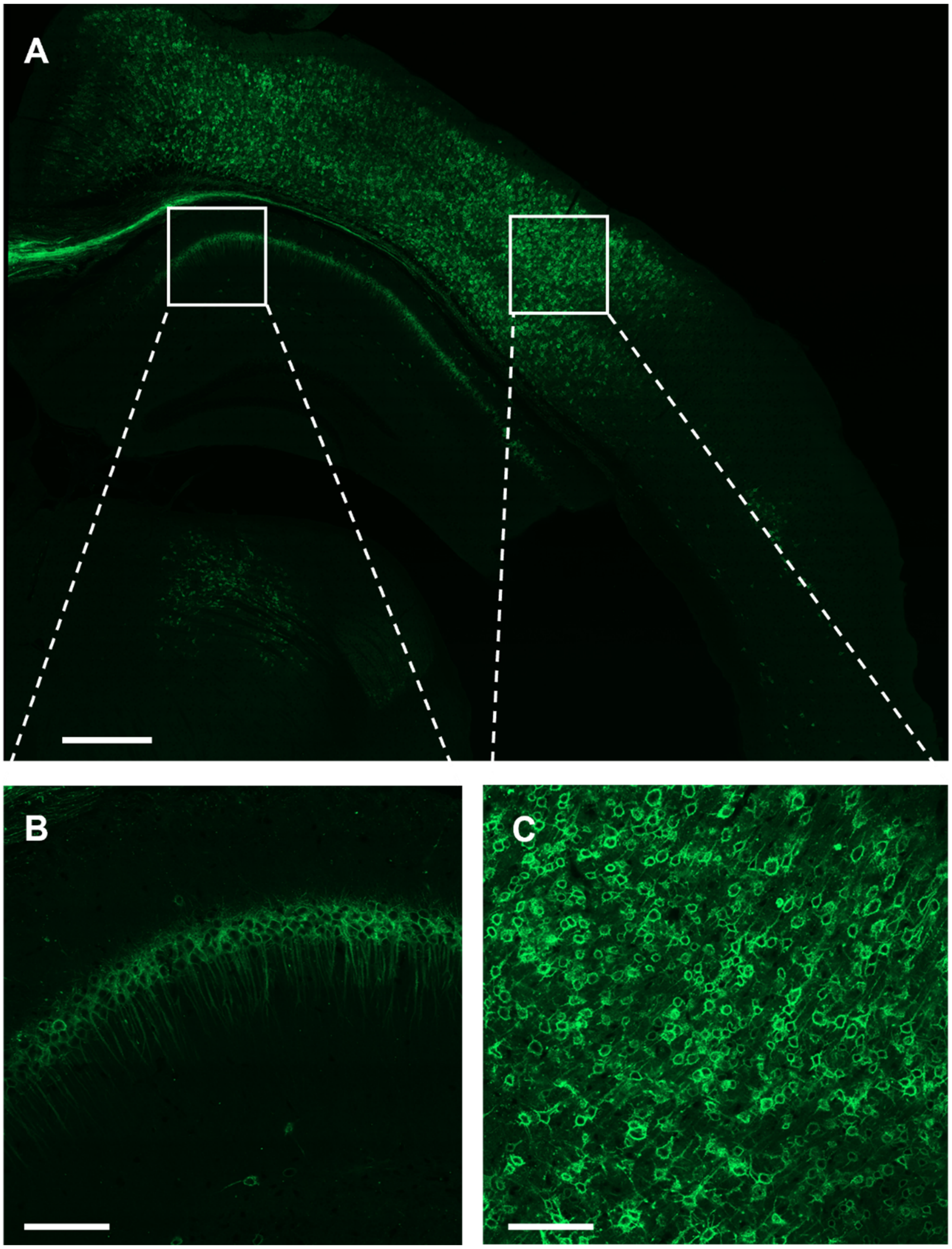

**Figure S6. Confocal imaging in formaldehyde-fixed brain slices expressing Vega.** **(A)** Confocal images of fixed slices showing excellent expression and localization of Vega. AAV encoding Vega was injected into V1 cortex. Scale bar, 500 μm. **(B-C)** Zoomed-in figure of hippocampus (B) and cortex (C). Scale bars, 100 μm**.**

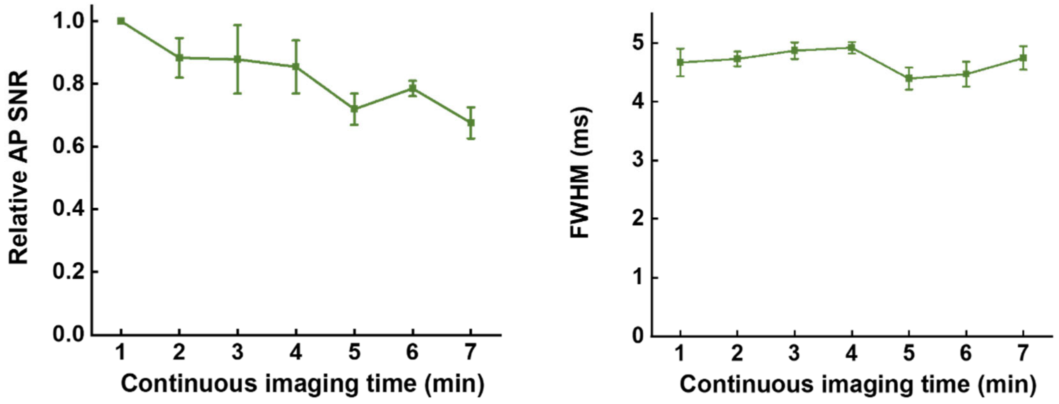

**Figure S7. Statistics of APs of Ace-mNeon2 during long-term continuous imaging in acute brain slices.** Normalized SNR (left) and average FWHM (right) of APs recorded by Ace-mNeon2 in each 1-min imaging session. n = 4 slice recorded from 2 mice. Error bars represent s.e.m.

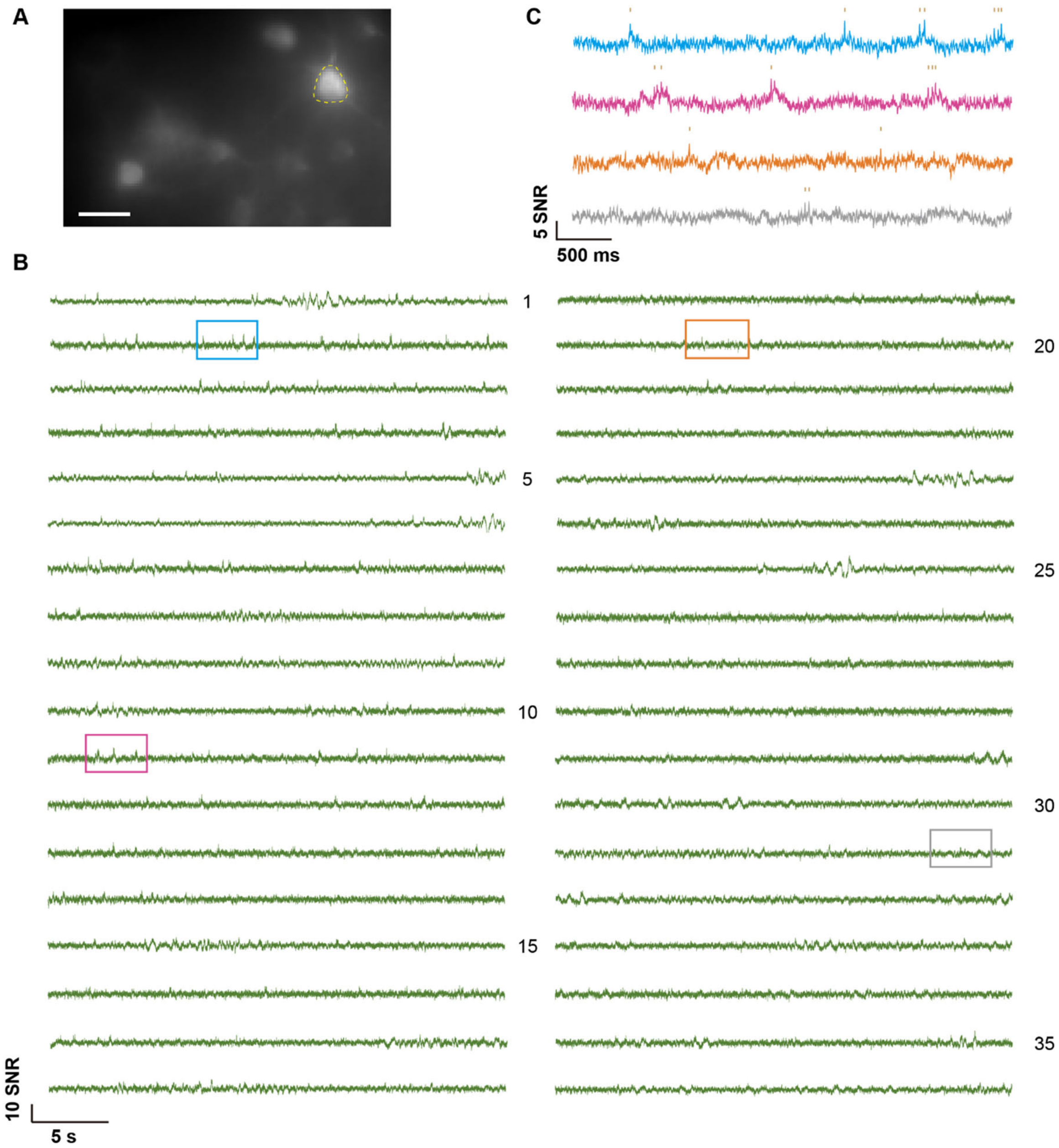

**Figure S8. 18-min voltage recording of Vega1.2-ST, expressing in V1 layer 2/3 pyramidal neurons (PNs).** Imaging were conducted in 30-sec sessions with 30sec-1minute intermittency. **(A)** Projection image showing L2/3 PNs in V1 expressing Vega1.2-ST *in vivo* with the target neuronal ROI highlighted in yellow. **(B)** Full 18-min fluorescence traces from the neuronal ROI in (A). **(C)** 4-sec trace snippets showing the voltage recording from the colored box area in (B). Scale bar = 25μm.

### Supplementary Methods

#### Materials and reagents

The reagents used in this study are summarized in table S4. All animal maintenance and experimental procedures were conducted according to institutional and ethical guidelines for animal welfare, and all animal studies were approved by Institutional Animal Care and Use Committee of Peking University and the Institutional Animal Care and Use Committee (IACUC) of Westlake University, Hangzhou (animal protocol #KP-19-044) and . C57BL/6J mice were used in this study regardless of sex.

#### HEK293T cell culture and transfection

Human embryonic kidney (HEK) 293T cells were incubated in Dulbecco's modified eagle medium (DMEM, Gibco™) supplemented with 10% v/v fetal bovine serum (FBS, Gibco™) in an incubator at 37 °C with 5% $CO_2$. For experiments, HEK293T cells were seeded in a 24-well plate. At 70-90% confluency, cells in each well were transfected by using 500 ng plasmid and 1 μL Lipofectamine™ 2000 Reagent mixed in Opti-MEM™ Medium for 4 h. After that, cells were digested by Trypsin-EDTA (0.25%, Gibco™), reseeded on Matrigel® Matrix-coated 14-mm glass coverslips, and then cultured in cell medium for 24 h before labeling with fluorophore.

#### Neonatal rat neuron culture and transfection

For primary rat hippocampus neuron culture, 14-mm glass coverslips were pre-coated with poly-D-lysine (Sigma) solution and Laminin Mouse Protein (Gibco™) solution at 37 °C within 2 days. After that, the coverslips were washed twice with $ddH_2O$ and let dry at room temperature. To isolate neurons, neonatal Sprague-Dawley rats' heads were cut off with scissors. Then, the brain was isolated from the skull and put into the ice-chilled dissection solution (DMEM with high glucose and penicillin-streptomycin antibiotics). Next, the hippocampi were separated from brains under a dissection scope, cut into small pieces, and incubated with Trypsin-EDTA (0.25%) for 15 min at 37°C. Thereafter, the trypsin was gently replaced with DMEM containing 10% FBS. After being repeatedly pipetted for 30 second and incubated on ice for 5 min, the tissue fragments were sedimented at the bottom of the centrifuge tube. The supernatant was collected and diluted with neural culture medium (Neurobasal™ medium supplemented with B-27™ supplement, GlutaMAX™ supplement, and penicillin-streptomycin) to a final cell density of $6\times10^4$

cells/mL. 1 mL cell suspension was added to each 24-well containing one coverslip. Every four days, half of the neural culture medium was replaced with fresh medium.

For the transfection experiments, cultured neurons were transfected at DIV8 (8 days in vitro). Neurons for each 24-well were treated with a mixture of 0.75-1 μg plasmids and 1 μL Lipofectamine™ LTX for 40 min. After 4-6 days, the transfected neurons were labeled and imaged.

**Imaging apparatus, wide-field microscopy and confocal microscopy**

Wide-field fluorescence imaging experiments in HEK293T cells, rat cultured neurons and mouse pancreatic islets were performed with an inverted fluorescence microscope (Nikon-TiE) equipped with a 40× 1.3 NA oil immersion objective lens, Coherent OBIS 488 nm laser line, and scientific CMOS camera (Hamamatsu ORCA-Flash 4.0 v2). The microscope, lasers, and cameras were controlled by LabVIEW (National Instruments, 15.0 version) software.

Wide-field fluorescence imaging experiments in brain slice were performed with an upright lab-build fluorescence microscope equipped with a Nikon 25× 1.1 NA water immersion objective lens with an f = 60 mm tube lens from ThorLab, SPECTRA X Light Engine Cyan channel (Bandpass 475/28) and scientific CMOS camera (Hamamatsu ORCA-Fusion). The cameras were controlled by HCImageLive software from Hamamatsu. The LED was controlled by Lumencor Light Engine software from Lumencor.

Table S3 provides a summary of the filters and dichroic mirrors utilized for the fluorescence indicators in this work. Images of HEK293T cells, rat cultured neurons and mouse pancreatic islets were captured using a camera on 2×2 pixel binning and a 100-ms exposure duration, and images of brain slices were captured using a camera on 1×1 pixel binning and a 2.5-ms exposure duration. Images were analyzed using MaTLAB (version R2022a) software.

Confocal fluorescence imaging experiments in brain slice were performed with an inverted confocal-laser scanning microscope (ZEISS LSM 900) equipped with a 63× 1.4 NA oil immersion objective lens and an inbuilt 488nm laser with a power of 4.5% of the maximum power. The microscope, lasers, and cameras were controlled by ZEN (ZEISS, 3.6 version) software. Images were analyzed using ZEN (ZEISS, 3.6 version) software.

**Electrophysiology**

For single-cell electrophysiological recording, cultured neurons were incubated in Tyrode's buffer containing 20 μM gabazine, 10 μM NBQX and 25 μM APV (Tyrode’s buffer containing 50 nM 2-APB for HEK293T cells). The electrophysiological experiments were conducted at room temperature. Borosilicate glass electrodes (Sutter) were pulled to a tip resistance of 2.5-5 MΩ. The internal solution containing 125 mM potassium gluconate, 8 mM NaCl, 0.6 mM $MgCl_2$, 0.1 mM $CaCl_2$, 1 mM EGTA, 10 mM HEPES, 4 mM Mg-ATP, 0.4 mM GTP·$Na_2$ (pH 7.3) was adjusted to 295 mOsm/kg with 1 M sucrose and injected into the glass electrode. The glass electrode's position was adjusted by a Sutter MP285 micro-manipulator. An Axopatch 200B (Axon Instruments) amplifier was utilized to clamp the cells. Membrane potential data recorded from the amplifier was filtered by an internal 5 kHz Bessel filter and digitalized at 9681.48 Hz with a National Instruments PCIe-6353 data acquisition (DAQ) board.

**Simultaneous patch clamp and optical recording in cultured cells**

The membrane potential was controlled by whole-cell patch clamp. To characterize the dynamic range of voltage indicators in HEK293T cells, the membrane potential was stepped from -100 mV to 100 mV in 20 mV increments, lasting 500 ms for each step. Meanwhile, the fluorescence images were recorded at 1058 Hz.

Neuronal single APs were stimulated by injecting 100–400 pA current for 10 ms into the cultured neurons. Fluorescent signals were acquired at 484 Hz camera frame rate with 2-by-2 binning.

**Intracerebroventricular injection and acute slice measurements**

The AAV vector expressing Vega: AAV2/9-hsyn-Vega was custom-produced by Chinese Institute for Brain Research. For one 6- to 7-week-old mouse (without regard to sex), 200 nL of AAV2/9-hsyn-Vega ($1.4 \times 10^{13}$ genome copies/mL) was injected into the thalamus. The coordinate for intracerebroventricular injection (in millimeters from Bregma: anteroposterior and mediolateral) was 1.05 and 0.3 and dorsoventral of 5.3 mm.

Acute slices were prepared from 9- to 11-week-old mice (at least 3 weeks after AAV injection). Mouse was deeply anesthetized via isoflurane inhalation and rapidly decapitated. The brain was dissected from the skull and placed in ice-cold artificial cerebrospinal fluid (ACSF) containing 26 mM $NaHCO_3$, 1.25 mM $NaH_2PO_4$, 125 mM NaCl, 2.5 mM KCl, 2 mM $CaCl_2$, 1 mM $MgCl_2$, 2 mM KCl, and 20 mM glucose (295 mosmol/kg; pH 7.3 to 7.4) and saturated with Carbogen (95% $O_2$ and 5% $CO_2$). The brain was sliced into 200-μm sections with a Leica VT1200s vibratome. Slices were incubated for 45 min at 34.5°C in ACSF and maintained in ACSF (34.5°C). ACSF was continuously bubbled with Carbogen for the duration of the preparation and subsequent experiment.

For fluorescence imaging and electrophysiology recoding, slices were placed on a custom-built chamber and held by a platinum harp net stretched across. Carbogen-bubbled ACSF was perfused at a rate of 1 mL/min with a longer peristaltic pump. Fluorescence images were captured at a frame rate of 400 Hz, under 488-nm illumination (1 W/cm$^2$).

**Isolation of mouse pancreatic islets and infection of Vega**

Islets of Langerhans were isolated from wild-type C57/6j mice. After isolation, the islets were cultured overnight in RPMI 1640 medium containing 10% fetal bovine serum, 8 mM d-glucose, penicillin (100 U/mL), and streptomycin (100 mg/mL) for overnight culture at 37°C in a 5% $CO_2$-humidified air atmosphere. The islets were infected with adenoviruses pADV-prip2-Vega by 1.5-hour exposure in 100 μL of culture medium, with approximately 1 × $10^{10}$ plaque forming units per islet, followed by addition of regular medium and further culture for 36 to 48 hours before use.

**Voltage imaging in mouse pancreatic islets**

For imaging with Vega, mouse pancreatic islets were illuminated with 488 nm laser at 1.5 W/cm$^2$, and continuously imaged for 180 s at a camera frame rate of 200 Hz for five rounds. The samples were kept at 37°C on the microscope stage during imaging. The mouse islets were plated on poly-l-lysine–coated coverslips in the glass bottom dish, and were treated with KRBB solution [125 mM NaCl, 5.9 mM KCl, 2.56 mM $CaCl_2$, 1.2 mM $MgCl_2$, 1 mM l-glutamine, 25 mM Hepes, and 0.1% bovine serum albumin (pH 7.4)] containing 3 mM d-glucose for 15 minutes to silence APs before imaging. After 3-minute imaging, d-glucose was added to 13 mM manually, and performed the next four rounds of imaging continuously. All displayed traces of Vega in mouse pancreatic islets did not undergo photobleaching correction.

**AAV injection and surgery in primary visual cortex mice**

Briefly, eight-week-old mice were anesthetized with 4% isoflurane for induction and maintained under 1.5–2% isoflurane throughout the procedure, with body temperature maintained on a 37 °C heating pad. The mice were positioned in a stereotaxic apparatus (RWD Life Science, USA). Scalp hair was removed using depilatory cream, and the eyes were protected with erythromycin ophthalmic ointment. A craniotomy approximately 3 mm in diameter was performed over the visual cortex at coordinates anteroposterior (AP): –3.4 mm and mediolateral (ML): 2.1 mm. For viral delivery, AAV2/9-CAG-DIO-Vega1.2ST (final titer ~1×$10^{13}$ vg/mL) and AAV2/9-CaMKIIa-Cre (final titer 1×1×$10^{10}$ vg/mL) were mixed. A volume of 100 nL of this mixture was injected into the primary visual cortex at AP: –2.5 mm, ML: 2.5 mm, and depth: 200 μm below the cortical surface. The injection was delivered at a rate of 50 nL/min. A total of four to five injections were administered in the left visual cortex, spaced 300 μm apart, at coordinates AP: –2.5 mm and ML: 2.5 ± 1 mm. Immediately after the viral injection, a coverslip (3 mm in diameter, 0.1 mm thick; Luoyang GULUO Glass) was placed over the exposed dura mater. To seal the imaging window, Kwik-Sil adhesive (World Precision Instruments) was applied around the edges of the coverslip. The entire skull was then covered with dental cement (C&B Metabond, Parkell), and a custom titanium headplate was affixed atop the cement. Once the cement had hardened, denture base resin was applied to reinforce the window assembly. The mice were allowed to recover for at least 3–4 days before undergoing further experiments.

**Wide-field one-photon imaging in awake mice in primary visual cortex**

After 3–4 weeks following viral infection and window implantation, the mice were prepared for imaging experiments. Prior to image acquisition, the mice were anesthetized with 2% isoflurane and head-fixed onto a custom imaging stage. Wide-field imaging was performed using an epifluorescence microscope (Olympus BX51WI) equipped with a ×20 water-immersion objective lens (NA 0.95) and an sCMOS camera (OrcaFlash4.0v2, Hamamatsu). Epi-illumination was provided by a 473 nm laser (Changchun New Industries Optoelectronics Tech), adjusted to power densities between 5 W/cm² and 10 W/cm². Neurons located at depths of 130–200 μm below the pia mater were recorded. To ensure efficient image transfer and enable prolonged recording sessions, we conducted consecutive imaging trials. Specifically, 19,140 frames were acquired at a rate of 638 Hz during each 30 s continuous recording session (image dimensions: 80–120 × 80 pixels, with 4 × 4 pixel binning), followed by a timed 30-s dark period without imaging.

**Data analysis**

Both electrical data and fluorescence images were analyzed by the custom MATLAB software (MathWorks, version R2022a). For each cell, fluorescence intensities were taken from the mean pixel values of a manually created ROI around the soma. After camera bias removal (100 and 400 for 1-by-1 and 2-by-2 binning, respectively), the raw optical traces of spontaneous electrical activities averaged from ROIs were corrected with background subtraction and photobleaching removal. The trend lines were generated by a smooth function using a 2.5-s (999 consecutive data points) moving average to the background-subtracted data. The photobleaching of the traces was removed by normalizing to the corresponding trend lines unless otherwise noted. Statistical analyses were performed using Excel (Microsoft Excel 2019) and Origin (version 2024b).

**Data Availability Statement**

Data presented in this study are provided in Supplementary Tables.
